# Motor Resonance of Musical Emotion: A Machine Learning Approach to EEG Decoding During Expressive Music Performance

**DOI:** 10.64898/2026.06.04.730044

**Authors:** Alice Mado Proverbio, Miloš Milovanović

**Author notes:** Correspondence, AMP, Department of Psychology of University of Mila-no-Bicocca, Piazza dell’Ateneo Nuovo 1, 20162 Milan, Italy.

## Abstract

Understanding the neural dynamics underlying expressive musical performance remains a major challenge at the intersection of neuroscience, music cognition, and computational modeling. While EEG studies of emotion have largely focused on passive exposure to affective stimuli, comparatively little research has examined oscillatory brain activity during active musical expression. The present single-subject study investigated whether band-limited EEG activity recorded during expressive piano performance by a professional concert pianist contains sufficient discriminative structure to support supervised multi-class classification of musically defined emotional categories.

**Methods:** EEG was recorded from 128 scalp sites while a professional concert pianist performed emotionally characterized excerpts from Bach, Beethoven, and Chopin in a continuous naturalistic session. Musical excerpts had been previously categorized and perceptually validated according to emotional valence, tempo, energy/arousal, and tonal structure. From the continuous EEG recording, 180 non-overlapping 2-second artifact-free segments were extracted, yielding 30 segments for each emotional category. Mean spectral power was computed within theta (3.5-7.5 Hz), alpha (7.5-12.5 Hz), and high-beta (24-30 Hz) frequency bands across selected centro-parietal and posterior electrodes, resulting in 24 EEG-derived features per segment. Linear Support Vector Machine, Random Forest, and Gradient Boosting classifiers were evaluated using an 80/20 train-test split combined with 5-fold cross-validation.

**Results:** EEG-only classification achieved above-chance performance across models, with Random Forest yielding the highest accuracy (0.42), macro F1-score (0.414), and Cohen’s κ (0.30), exceeding the theoretical chance level of 0.167. Feature importance analysis revealed distributed contributions across theta, alpha, and high-beta oscillatory activity, particularly over parietal and occipital regions, without evidence for a single dominant neural marker. Inclusion of an additional binary arousal-related feature substantially improved Random Forest performance (accuracy = 0.58; macro F1 = 0.579; κ = 0.50), indicating that arousal organization contributed strongly to category separability within the classification framework.

**Conclusions:** These findings suggest that oscillatory EEG activity accompanying expressive musical action contains measurable statistical structure associated with emotionally differentiated performance states. Rather than identifying discrete neural correlates of emotion, the present results provide a computational characterization of distributed oscillatory dynamics emerging during expressive motor-acoustic interaction, extending affective EEG research beyond passive perception paradigms toward ecologically grounded musical performance contexts.

## 1. Introduction

This study provides an exploratory computational analysis of oscillatory EEG activity during live expressive piano performance, testing whether frequency-domain features can support classification of predefined emotional categories under ecologically valid sensorimotor conditions. Musical performance requires continuous integration of motor execution, auditory feedback, temporal control, and expressive intention. In piano performance, systematic variations in dynamics, articulation, and temporal structure generate acoustic signatures associated with emotional categories [1]. Prior work links emotion to measurable performance parameters such as intensity, note density, gesture rate, and temporal structure [2], while removal of expressive acoustic cues significantly impairs emotion recognition [3,4]. Electroencephalography has been widely used in affective neuroscience to study neural correlates of emotional processing [5,6], with a shift over the past decades from ERP-based descriptions to computational decoding of affective states from oscillatory EEG features. This transition reflects advances in machine learning enabling classification from multivariate frequency-domain representations. Standard EEG decoding pipelines involve preprocessing, feature extraction, and supervised classification, with band-limited spectral power remaining a central feature due to its interpretability and robustness [7–9]. Benchmark datasets such as DEAP [10], SEED [11], and DREAMER [12] have shaped this field. These datasets typically involve multi-subject recordings during exposure to emotionally evocative audiovisual stimuli, with affect annotated along valence and arousal dimensions. Valence reflects hedonic quality, whereas arousal indexes physiological activation; arousal is associated with widespread modulation of cortical oscillatory activity [13]. Across these datasets, EEG is segmented into short windows, transformed into the frequency domain, and summarized using bandpower features. Feature extraction commonly relies on spectral power in canonical frequency bands (delta, theta, alpha, beta, gamma), complemented by connectivity, asymmetry, or entropy-based metrics. Among these, bandpower remains most widely used due to its efficiency and interpretability [14–16]. Classification approaches include SVMs, k-NN, Random Forests, Logistic Regression, and more recently CNNs and RNNs [14,17,18]. Classical models remain competitive in low-data regimes due to robustness and interpretability, whereas deep learning typically requires larger datasets and stronger regularization [8,14]. A key methodological issue is subject-dependent versus subject-independent decoding, with consistently higher performance in within-subject settings and persistent challenges in cross-subject generalization due to inter-individual variability and anatomical differences [8,11,19,20]. Despite progress in EEG-based affective decoding, most studies rely on passive perception paradigms, with emotional states elicited through external audiovisual stimuli rather than motor production, limiting ecological validity for action-based affect.

Within embodied cognition frameworks, expressive meaning emerges from coupling between motor planning, execution, and auditory feedback [21–23]. Oscillatory EEG activity during performance may therefore index sensorimotor coordination underlying expression. Theta (4-8 Hz), alpha (8-13 Hz), and beta (13-30 Hz) bands have been linked to temporal organization, attentional modulation, and motor control, respectively. Beta desynchronization accompanies movement execution [24,25], alpha reflects inhibitory control and resource allocation [26,27], and theta supports temporal and cognitive control [28,29], jointly enabling integration of motor and auditory processes in music performance [30,31]. Interestingly, Zamm et al. [32] demonstrated that motor-related EEG can be recorded during naturalistic musical performance using wireless EEG synchronized with MIDI, enabling precise neural-behavioral alignment. Virgilio et al. [33] further showed that motor patterns across hand, foot, and tongue movements are decodable above chance, supporting somatotopic organization and multiclass BCI feasibility. Machine learning extends EEG analysis beyond univariate approaches by enabling decoding of distributed neural patterns [8,14,7]. Classical models such as SVMs, Random Forests, and Gradient Boosting remain widely used due to robustness in small-sample settings and interpretability [34,9], while deep architectures (CNNs, RNNs) can learn hierarchical representations but require larger datasets and are prone to overfitting [35,18]. Consequently, feature-based approaches remain competitive in constrained datasets. Model evaluation is typically performed using subject-dependent cross-validation schemes [20,36].

In this study, EEG was recorded from a professional concert pianist performing emotionally characterized excerpts from Bach, Beethoven, and Chopin. Frequency-domain features were extracted from centro-parietal and parieto-occipital electrodes and used in supervised classifiers including Linear SVM, Random Forest, and Gradient Boosting. The study (i) investigates emotional decoding during live performance, (ii) quantifies band-specific contributions of oscillatory power, and (iii) evaluates interpretable multiclass classification models in an ecologically valid EEG setting. EEG was recorded during continuous expressive piano performance and subsequently transformed into frequency-domain representations suitable for computational analysis. The methodological design aimed to preserve ecological validity while enabling structured classification across predefined musical emotion categories.

## 2. Materials and Methods

The analysis pipeline followed a feature-based supervised learning approach. Continuous EEG recordings were segmented into temporally defined windows aligned with emotionally labeled musical excerpts. Spectral power was computed within predefined frequency bands and across selected scalp electrodes. These band-limited power values served as input features for multi-class classification using established machine learning models. Given the single-participant design, methodological decisions emphasized reproducibility, statistical stability, and transparent feature-level analysis. Classical supervised learning algorithms were investigated due to their robustness in moderate-dimensional feature spaces and their capacity to estimate relative feature contributions within the classification task.

### 2.1. Participants

The participant is a 30-year-old right-handed concert pianist of international distinction who commenced formal piano training at the age of four and graduated with highest honors from a world-renowned conservatory. Over the course of his career, he has cultivated extensive international concert experience, performing at eminent venues and in collaboration with prestigious musical institutions worldwide, including New York’s Carnegie Hall and Nagoya Philharmonic Orchestra. Accordingly, he possessed the technical command and interpretative sophistication necessary to execute the experimental repertoire with both virtuosity and expressive nuance. He reported normal auditory and visual function and no history of neurological, psychiatric, motor, or cognitive disorders. Experimental sessions were conducted in an anechoic chamber using a Yamaha P-225B digital piano configured with the default grand piano timbre. Audio performances were recorded in WAV format while EEG and behavioral data were acquired concurrently. The experimental repertoire comprised six works, amounting to approximately 27 minutes of performance. All pieces were performed from memory and were informed by interpretative frameworks derived from recent commercial recordings as well as the pianist’s established touring repertoire. Performances were rendered with stylistic coherence, interpretative naturalness, and a high degree of expressive consistency following a brief period of focused preparatory concentration.

### 2.2. Stimulus material

Within the performed repertoire, 110 musically meaningful phrases lasting about 10-11 sec were identified as representing different emotional categories and musical styles. Musical excerpts were taken from: J. S.□Bach’s Die Kunst der Fuge (BWV□1080), specifically Contrapunctus I, VII, IX, and XI (Baroque period, characterized by intricate counterpoint, harmonic clarity, and affective expressiveness); Ludwig van Beethoven’s Piano Sonata Op.□110, Adagio (transition from Classical to early Romantic style, marked by formal innovation, dynamic contrasts, and heightened emotional depth); and Frédéric Chopin’s Ballade No.□1 in G minor, Op.□23 (romantic period, known for lyrical phrasing, harmonic richness, and introspective expressivity). By incorporating excerpts from stylistically distinct yet universally recognized composers, we aimed to assess whether structural and perceptual properties of music—such as tonality, tempo, complexity, and energetic content—generate distinguishable EEG signatures related to audiomotor processing, irrespective of composer identity. Musical categories were established through structural analysis of the excerpts and subsequently validated perceptually using audio-only presentation in a cohort of 20 trained musicians and independent listeners, who showed reliable agreement across categories [2]. In parallel, objective differences in acoustic and performance-related parameters, including intensity (dBFS), note density, gesture rate, and key, systematically mapped onto the predefined emotional categories (Table 1).

**Table 1.** Emotional categories used for classification of 110 musical excerpts. While the structural differences were objective, the emotional label reflected a psychological sensation.

| Emotional valence | Emotional feeling | Key | Tempo | Energy/Arousal |
| --- | --- | --- | --- | --- |
| Positive | Peacefulness (PEA) | Major | Slow | Little |
| Positive | Joy (JOY) | Major | Fast | Considerable |
| Negative | Melancholy (MEL) | Minor | Slow | Little |
| Negative | Argumentative (ARG) | Minor | Fast | Some |
| Negative | Tension (TNS) | Minor | — | Considerable |
| — | Power (POW) | — | — | Strong |

Participants’ emotion judgments were analyzed within a supervised multi-class classification framework, allowing systematic quantification of agreement and misclassification patterns relative to predefined target categories. To improve classification reliability, auditory excerpts with low recognizability were excluded, resulting in a reduced set of 56 musical fragments. The discarded fragments showed anyhow an above chance accuracy rate of 47% (chance threshold being 16.67%). Overall classification performance yielded mean values of Precision = 0.61, Recall = 0.60, and F1-score = 0.60 (see Fig. 1), which exceeded our expectations. Subsequently, 180 labeled EEG segments were obtained by extracting 30 artifact-free, non-overlapping 2-second epochs for each emotional category and composer, based on the previously validated musical categorization. The segments were not temporally continuous; rather, they were sampled from distinct time points distributed across the full duration of the performance recording. This design yielded a balanced multi-class dataset with equal representation across all six emotional conditions.

**Figure 1.**
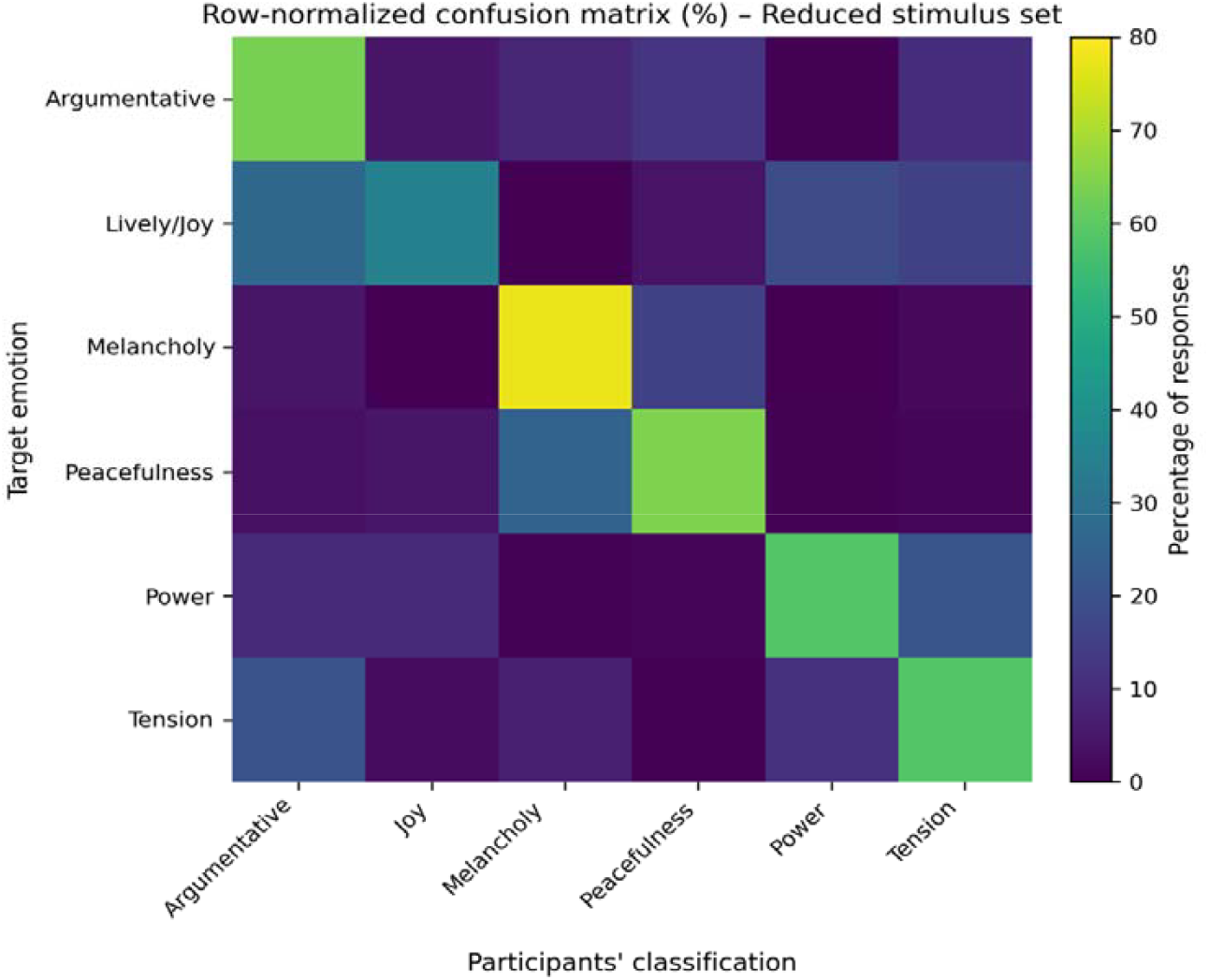
Row-normalized confusion matrix for the reduced stimulus set (56 musical fragments). Rows represent target emotion categories and columns represent participants’ classifications. Cell color indicates the percentage of responses assigned to each category within each target emotion.

### 2.3. EEG recordings and analysis

Scalp EEG was recorded from 128 Ag/AgCl electrodes arranged according to the 10-5 International System using ANT Neuro Waveguard high-density caps. EEG recordings were referenced online to the algebraically averaged mastoids. Electrode impedances were maintained below 5 kΩ throughout acquisition. Continuous EEG data were digitized at a sampling frequency of 512 Hz using the Cognitrace acquisition platform (ANT Software). EEG and EOG signals were amplified and band-pass filtered within a frequency range of 0.016-70 Hz. Artifact rejection was performed automatically prior to averaging, excluding epochs containing voltage fluctuations exceeding ±50 μV at any scalp electrode. Stimulus-locked EEG epochs were subsequently segmented and processed offline using the EEProbe analysis environment (ANT Software). While the continuous EEG recording encompassed the full duration of the performance EEG analyses focused on temporally defined segments corresponding to pre-defined emotionally categorized musical excerpts. To illustrate the structure of the recorded signals, a representative 2-second raw EEG segment is shown in Figure 2, separately for the 6 categories.

**Figure. 2.**
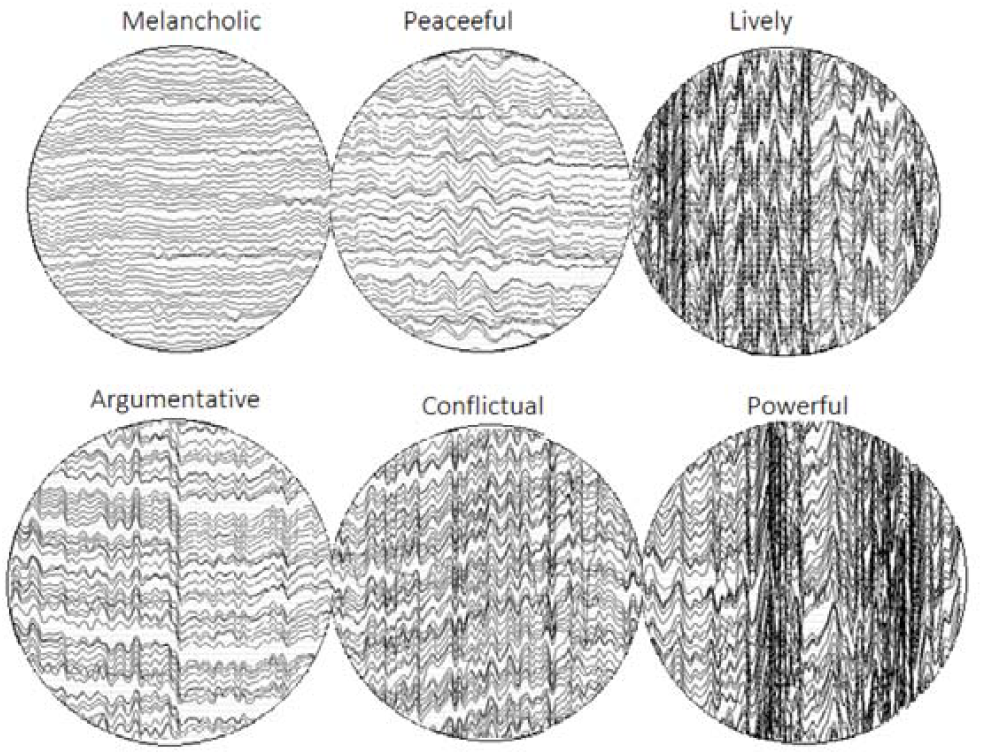
Representative 2-second raw EEG segment recorded during expressive piano performance of Bach

### 2.4 Feature Engineering: Preprocessing and Frequency-Domain Transformation

EEG signals recorded by each electrode were band-pass filtered between 0.016 Hz and 70 Hz. Automatic artifact rejection procedures were applied inside of the ASA software prior to spectral analysis. The continuous signal was segmented into non overlapping 2-second windows corresponding to the predefined emotional excerpts. For each 2-second segment, a Fast Fourier Transform (FFT) was computed for the initial set of 18 electrodes to obtain the frequency spectrum. Mean spectral power was calculated within predefined frequency bands: Delta (0.5-3.5 Hz), Theta (3.5-7.5 Hz), Alpha (7.5-12.5 Hz), Beta-low (12.5-18 Hz), Beta-mid (18-24 Hz), Beta-high (24-30 Hz), Gamma (35-45 Hz) and served as the primary frequency-domain descriptor. The initial dataset included information from a broader set of frontal, central, parietal, and occipital electrodes. Guided by neurophysiological assumptions [26,37] and considering the task at hand, the initial electrode set was reduced to the following eight posteriori and rolandic channels: O1, O2, PO3, PO4, P3, P4, CP1, CP2. In a similar manner, certain frequency bands were removed from consideration. The gamma band was excluded because no significant task-related modulation was observed, and high-frequency activity in scalp EEG is typically weak, transient, and more susceptible to muscle contamination. The delta band was excluded due to its susceptibility to slow drifts and non-neural artifacts [38,39]. Low and mid-beta ranges were excluded on the basis that continuous motor execution. during performance may produce sustained event-related desynchronization, potentially reducing their discriminative value [25,26]. The final frequency bands retained for classification were: Theta (3.5-7.5 Hz), Alpha (7.5-12.5 Hz), Beta-high (24-30 Hz). The final set of features consisted of 24 band-limited power values per segment, corresponding to three frequency hands (theta, alpha and high beta)extracted from eight selected electrodes In addition to EEG-derived features, an artificial binary feature representing arousal was introduced for comparative analysis. This feature was assigned as follows: 0 → MEL, PEA, ARG; 1 → JOY, POW, TNS. This feature represents a coarse cortical arousal-based categorization of emotional conditions under investigation. Since it is derived from the emotional labels themselves, it introduces partial label information into the feature space. For this reason, analyses were conducted both with and without inclusion of this feature in order to evaluate its influence on classification performance.

### 2.5. Machine Learning Pipeline: Classification Framework

Multi-class classification was performed across the six emotional categories (MEL, PEA, ARG, JOY, POW, TNS). The dataset was randomly shuffled prior to model training to avoid ordering effects. A supervised learning framework was implemented using the following classifiers:

1. Linear Support Vector Machine (SVM)
2. Random Forest
3. Gradient Boosting

These models were selected due to their suitability for moderate-dimensional feature spaces and their established use in EEG-based decoding research.

#### 2.5.1 Linear Support Vector Machine (SVM)

It is a margin-based classifier that seeks to identify a hyperplane separating classes by maximizing the margin between decision boundaries [40]. In the multi-class setting, a one-vs-rest strategy was employed. Linear SVMs are particularly well-suited for structured but moderately sized feature spaces, where they provide stable decision boundaries and reduced risk of overfitting compared to highly flexible nonlinear models [36, 41]. Given the limited dataset size (180 segments), the Linear SVM served as a strong baseline classifier. Hyper-parameter tuning for the Linear SVM was performed using GridSearchCV with 5-fold stratified cross-validation. Model selection was based on macro F1-score to ensure balanced multi-class evaluation. The regularization parameter C was explored over the range {0.1, 0.5, 1, 2, 3, 5, 10, 15, 20}. For the EEG-only feature set, the optimal configuration corresponded to C = 10. When the arousal feature was included, the optimal value shifted to C = 3.

#### 2.5.2. Random Forest

It is an ensemble learning method that constructs multiple decision trees using bootstrapped subsets of the data and random subsets of features. Final predictions are obtained through majority voting across trees. This ensemble approach reduces variance and improves generalization relative to single decision trees. Random Forest models are capable of capturing nonlinear relationships and interactions between features, which is advantageous in EEG-based classification where oscillatory bandpower features may interact across electrodes and frequency bands. Additionally, Random Forest provides intrinsic feature importance estimates, supporting interpretability of model decisions [14].

Hyper-parameter tuning was performed using GridSearchCV with 5-fold stratified cross-validation. Model selection was based on macro F1-score to ensure balanced multi-class evaluation. The search space included variations in the number of trees (n estimators ∈ {25, 50, 100, 300}), maximum tree depth (max depth ∈ {None, 5, 10, 15}), minimum samples required for node splitting (min samples split ∈ {2, 5, 10}), minimum samples per leaf (min samples leaf ∈ {1, 2, 4}), and feature subsampling strategy (max features ∈ {sqrt, log2, 0.3}). Class weights were set to balanced.

The optimal hyper-parameter configurations obtained via cross-validated grid search are summarized in Table 2.

**Table 2.**
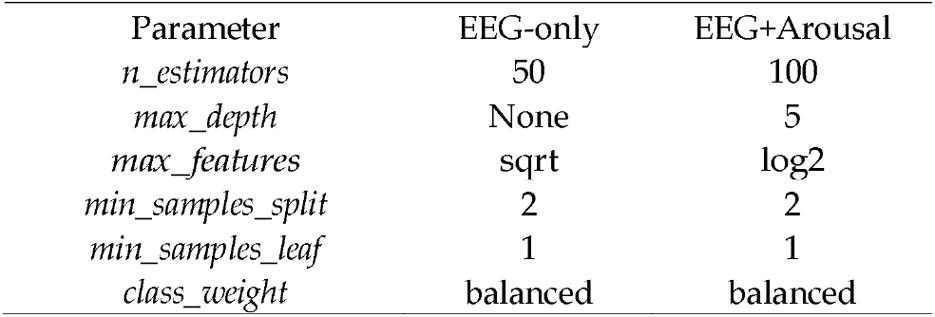
Optimal Random Forest hyper-parameters for EEG-only and EEG + arousal feature configurations.

| Parameter | EEG-only | EEG+Arousal |
| --- | --- | --- |
| *n_estimators* | 50 | 100 |
| *max_depth* | None | 5 |
| *max_features* | sqrt | log2 |
| *min_samples_split* | 2 | 2 |
| *min_samples_leaf* | 1 | 1 |
| *class_weight* | balanced | balanced |

#### 2.5.3 Gradient Boosting

It is an ensemble technique that builds decision trees sequentially, where each new tree is trained to correct errors made by the previous ensemble. Unlike Random Forest, which reduces variance through averaging, Gradient Boosting reduces bias by iterative error correction. While often achieving high predictive performance in structured datasets, Gradient Boosting can be sensitive to hyper-parameter selection and dataset size. Its inclusion allowed evaluation of a sequential ensemble approach relative to the parallel ensemble structure of Random Forest [42,43]. Hyper-parameter tuning for the Gradient Boosting classifier was performed using GridSearchCV with 5-fold stratified cross-validation. Model selection was based on macro F1-score to ensure balanced multi-class evaluation. The search space included variations in the number of boosting stages (n estimators ∈ {50, 100, 200}), learning rate (learning rate ∈ {0.01, 0.05, 0.1}), maximum tree depth (max depth ∈ {2, 3, 4}), minimum samples required for node splitting (min samples split ∈ {2, 5, 10}), minimum samples per leaf (min samples leaf ∈ {1, 2, 4}), and subsampling ratio (subsample ∈ {0.7, 0.9, 1.0}). The optimal hyper-parameter configurations for the EEG-only and EEG + arousal feature sets are summarized in Table 3.

**Table 3.** Optimal Gradient Boosting hyper-parameters for EEG-only and EEG + arousal feature configurations.

| Parameter | EEG-only | EEG+Arousal |
| --- | --- | --- |
| *n_estimators* | 50 | 50 |
| *learning_rate* | 0.05 | 0.05 |
| *max_depth* | 4 | 2 |
| *min_samples_split* | 2 | 10 |
| *min_samples_leaf* | 2 | 1 |
| *subsample* | 1.0 | 1.0 |

### 2.6 Training and evaluation

A 80/20 train-test stratified split was applied to partition the dataset. Within the training set, 5-fold cross-validation was used for model validation and hyper-parameter tuning. A fixed random state (random state = 42) was employed to ensure deterministic and reproducible results. Performance was evaluated using multi-class classification metrics.

For models permitting feature importance estimation (e.g. Random Forest), relative feature contributions were extracted to examine which frequency-band and electrode combinations most strongly influenced classification outcomes. These values reflect the statistical importance of features within the trained model and should be interpreted as indicators of feature relevance, not as evidence of specific neural sources.

## 3. Results

Performance was evaluated on the held-out test set, which represented 20% of all available data (36 segments). Classification accuracy, macro-averaged F1-score, and Cohen’s Kappa were used as primary evaluation metrics. Macro F1 was used to ensure equal weighting of all emotional categories, preventing dominant classes from disproportionately influencing the evaluation. Cohen’s kappa provided an agreement measure corrected for chance.

### 3.1. EEG-Only Classification

Classification performance using the 24 EEG-derived features (three frequency bands across eight electrodes) is summarized in Table 4. All evaluated models achieved accuracy above chance level (0.167), indicating that band-limited oscillatory features contained measurable discriminative structure across emotional categories.

**Table 4.** Classification performance using EEG-only features (24 band-specific spectral power features). Chance level accuracy for six classes is 0.167.

| Model | Accuracy | Macro F1-score | Cohen's $\kappa$ |
| --- | --- | --- | --- |
| Linear SVM | 0.39 | 0.299 | 0.267 |
| Random Forest | 0.42 | 0.414 | 0.300 |
| Gradient Boosting | 0.33 | 0.348 | 0.200 |

To further examine class-level prediction behavior, the confusion matrix for the Random Forest classifier (EEG-only features) is shown in Figure 3.

**Figure. 3.**
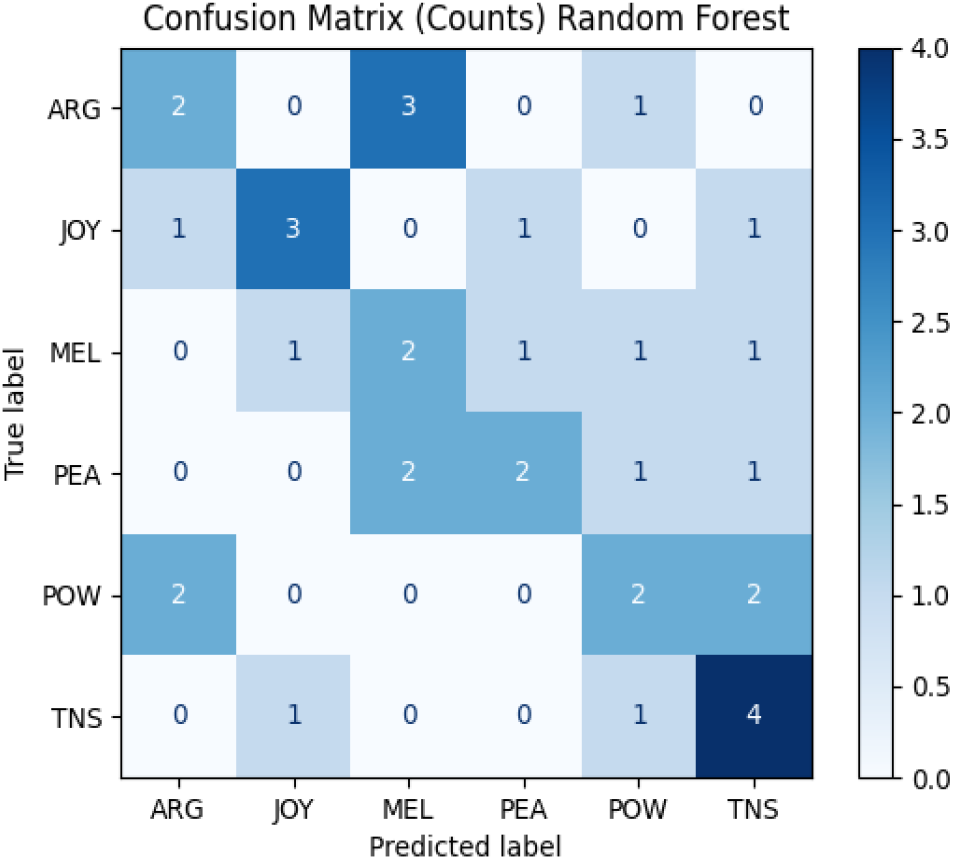
Confusion matrix for the Random Forest classifier using EEG-only features. Rows represent true labels; columns represent predicted labels.

Among the tested classifiers, Random Forest achieved the highest overall performance, suggesting that ensemble-based tree models were most effective in capturing nonlinear interactions within the frequency-domain feature space. Class-wise evaluation indicated relatively balanced performance across categories, with no class collapsing to zero recall. JOY (F1 = 0.55) and TNS (F1 = 0.53) demonstrated comparatively stronger separability, while MEL and POW exhibited more moderate discrimination.

Linear SVM demonstrated slightly lower overall performance. Although certain categories such as ARG and PEA were classified with relatively high recall, other categories showed reduced sensitivity, indicating less stable class separation compared to Random Forest. Gradient Boosting yielded the lowest EEG-only performance among the tested models. While individual categories such as JOY and POW demonstrated moderate precision, overall classification stability was reduced relative to the other approaches. Overall, EEG-only classification results indicate moderate but consistent separability between emotional conditions, reflecting partial discriminative structure within frequency-domain bandpower features.

### 3.2. Effect of Artificial Arousal Feature

To assess the influence of an explicit arousal-based grouping variable, a binary feature was introduced distinguishing lower-arousal (MEL, PEA, ARG) from higher-arousal (JOY, POW, TNS) emotions as determined by previous validation study [2]. Classification performance under this modified configuration is summarized in Table 5.

**Table 5.**
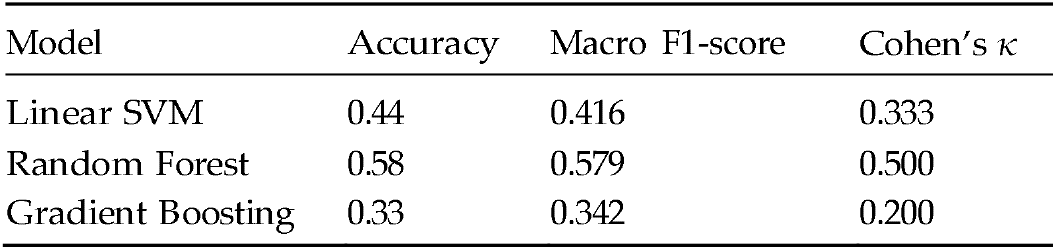
Classification performance including the artificial binary arousal feature.

The inclusion of the artificial arousal feature substantially improved Random Forest performance. Accuracy increased from 0.42 in the EEG-only case to 0.58, macro F1-score from 0.414 to 0.579, and Cohen’s κ from 0.30 to 0.50. Improvements were observed across multiple emotional categories, with ARG (F1 = 0.73), PEA (F1 = 0.67), and JOY (F1 = 0.60) demonstrating stronger discrimination relative to the EEG-only configuration.

Linear SVM exhibited a moderate performance increase following inclusion of the arousal feature, whereas Gradient Boosting did not show meaningful improvement. These results suggest that arousal is a meaningful dimension for distinguishing between emotional categories in the present dataset. However, an important methodological clarification is warranted regarding the interpretation of this feature. The binary arousal variable was constructed directly from the predefined emotional labels (0: MEL, PEA, ARG; 1: JOY, POW, TNS), meaning that it encodes category membership information rather than independently measured neural or behavioral data. Its inclusion therefore introduces partial label information into the feature space, which inflates classification performance in a manner that does not reflect neural decoding per se. This is not equivalent to data leakage in the strict sense—the feature is explicitly disclosed, its derivation is transparent, and analyses were conducted both with and without it—but its contribution to performance gains cannot be interpreted as evidence of additional oscillatory structure captured by the classifier. Rather, the arousal manipulation functions as an upper-bound reference condition, illustrating the degree to which arousal-level organization structures the six-class problem. The performance gap between EEG-only and EEG+arousal conditions (Δaccuracy = 0.16; Δκ = 0.20) quantifies how much discriminative information remains latent in the EEG features alone, and motivates future work on continuous, neurophysiologically-grounded arousal estimation. Consequently, EEG-only results provide the methodologically appropriate and interpretatively conservative basis for all conclusions regarding oscillatory contributions to emotional decoding during musical performance.

### 3.3. Feature Importance and Oscillatory Contributions

To further examine the relative contribution of individual oscillatory features, feature importance values were extracted from the EEG-only Random Forest model. The top-ranked features are presented in Table 6.

**Table 6.** Top 15 EEG feature importances derived from the Random Forest classifier using EEG-only features. Importance values reflect relative contribution to classification performance.

| Rank | Feature | Importance |
| --- | --- | --- |
| 1 | O2_high_beta | 0.0575 |
| 2 | P4_alpha | 0.0505 |
| 3 | P4_high_beta | 0.0497 |
| 4 | CP2_theta | 0.0496 |
| 5 | O2_theta | 0.0470 |
| 6 | O1_high_beta | 0.0463 |
| 7 | PO4_theta | 0.0438 |
| 8 | P4_theta | 0.0435 |
| 9 | O1_theta | 0.0435 |
| 10 | PO4_alpha | 0.0429 |
| 11 | CP2_high_beta | 0.0428 |
| 12 | P3_alpha | 0.0420 |
| 13 | O1_alpha | 0.0407 |
| 14 | PO4_high_beta | 0.0405 |
| 15 | PO3_alpha | 0.0400 |

The relatively narrow range of importance values suggests that classification relied on distributed multivariate patterns rather than a single dominant oscillatory marker. These importance measures reflect statistical contribution within the trained model and do not imply localization of specific neural generators. High-beta (24-30 Hz) features appeared frequently among the top-ranked contributors, including O2 high beta, P4 high beta, O1 high beta, CP2 high beta, and PO4 high beta. This repeated presence across occipital, parietal, and centro-parietal electrodes suggests that high-beta activity contributed consistently to class separation within the dataset. Theta-band features (3.5-7.5 Hz) were also strongly represented among influential predictors, including CP2 theta, O2 theta, PO4 theta, P4 theta, and O1 theta. The consistent contribution of theta across posterior and centro-parietal sites indicates that lower-frequency oscillatory dynamics also played a meaningful role in classification. Alpha-band features were present among top-ranked contributors (e.g., P4 alpha, PO4 alpha, P3 alpha, O1 alpha), though their relative importance values were slightly lower compared to high-beta and theta features. Overall, no single frequency band overwhelmingly dominated classification. Instead, theta, alpha, and high-beta bands contributed jointly to the model’s predictive structure. Spatially, important features were concentrated in parietal, parieto-occipital, and occipital electrodes (P3, P4, PO3, PO4, O1, O2), with additional contributions from centro-parietal sites (CP1, CP2). Frontal electrodes were not among the retained final feature set, consistent with the electrode reduction strategy previously described. The prominence of posterior and centro-parietal sites suggests that classification relied on distributed posterior oscillatory dynamics during expressive performance. Importantly, feature importance values reflect statistical relevance within the trained model and do not imply localization of specific neural generators.

## 4. Discussion

The present study examined whether oscillatory EEG activity recorded during expressive piano performance reflected structured motor-acoustic dynamics associated with predefined emotional categories. Using mean band-limited spectral power extracted from theta, alpha, and high-beta frequency bands across selected centro-parietal and posterior electrodes, moderate classification performance was achieved, far exceeding chance level (= 0.167) across models. EEG-only classification (accuracy = 0.42; macro F1 = 0.414) indicates that oscillatory dynamics during expressive motor production contain structured statistical information aligned with emotional categorization. However, the moderate magnitude of performance also reflects substantial overlap between categories. Emotional expression in music is continuous and multidimensional, and oscillatory brain activity reflects distributed neural coordination rather than discrete categorical states. Consequently, classification performance should be interpreted as evidence of partial separability within a complex neural feature space rather than as proof of distinct neural signatures for each emotional condition. Feature importance analysis of the EEG-only Random Forest model revealed that no single oscillatory feature dominated classification. Instead, contributions were distributed across high-beta, theta, and alpha bands, particularly over parietal, parieto-occipital, and occipital electrodes.

High-beta features appeared consistently among the most influential predictors, suggesting that fast oscillatory dynamics associated with motor coordination and sensorimotor engagement [26] strongly contributed to class separation. Theta-band features were also prominent, potentially reflecting integrative and timing-related processes inherent in structured musical execution [28]. Alpha-band contributions may relate to attentional regulation and functional coordination during continuous performance [27,44]. The present findings support the view that expressive musical performance is accompanied by distributed oscillatory coordination across multiple frequency bands, measurable at the scalp level and amenable to computational modeling [33].

The inclusion of the arousal feature substantially increased Random Forest performance (accuracy = 0.58; macro F1 = 0.579). These results suggest that arousal provides a highly informative organizational dimension that contributes strongly to emotional separability within the classification space. This finding is consistent with dimensional models of emotion, in which arousal represents a fundamental axis of affective organization associated with physiological activation and large-scale modulation of cortical dynamics [13,5]. Previous EEG-based affective decoding studies have similarly reported that arousal is often classified more reliably than valence, likely because variations in physiological activation are associated with broader and more robust oscillatory changes across cortical networks [10–11,15]. Within the present musical context, higher-arousal categories such as Joy, Power, and Tension were characterized by increased energetic intensity, faster temporal structure, and greater motor engagement, whereas lower-arousal conditions such as Peacefulness and Melancholy involved slower execution and reduced acoustic energy [2–4]. From an embodied sensorimotor perspective, these differences likely correspond to distinct patterns of movement coordination, attentional allocation, and auditory-motor integration, which may produce more globally separable oscillatory states during performance [45]. The improvement observed after inclusion of the arousal feature therefore supports the interpretation that expressive musical categories partly cluster according to underlying activation-related dynamics.

## 5. Conclusions

Overall, the results demonstrate that oscillatory bandpower features recorded during expressive motor production contain measurable statistical structure corresponding to predefined emotional categories [10,46]. Rather than identifying discrete neural generators, the findings characterize distributed oscillatory patterns associated with expressive performance. These patterns can be interpreted within an embodied framework in which emotional expressivity emerges from dynamic action-sound coupling during performance. These findings can be discussed in the light of the embodied hypothesis according to which emotional expressivity in musical performance emerges from dynamic patterns of action-sound coupling (musical gestures and corresponding sounds). Expressive differences between categories were previously shown to be associated with measurable performance parameters such as note density, acoustic energy, and gesture rate. High-arousal categories (e.g., Power, Tension, Joy) exhibited increased intensity and faster execution, whereas lower-arousal categories (e.g., Peacefulness, Melancholy) were characterized by reduced dynamic energy and slower temporal structure.

From an embodied perspective, such variations do not merely alter the acoustic signal but reflect distinct motor planning and execution dynamics. Increased note density and gesture speed require enhanced motor coordination and rapid sensorimotor integration, while lower-energy passages involve more sustained and temporally extended motor control. If emotional meaning is grounded in these structured motor-acoustic dynamics [2, 3, 4], corresponding differences in oscillatory brain activity would be expected to accompany expressive performance. The present EEG results are consistent with this view. Although classification performance was moderate, distributed oscillatory patterns across theta, alpha, and high-beta bands contained sufficient statistical structure to differentiate expressive categories above chance level. This suggests that neural activity recorded during performance reflects, at least in part, the organized motor-acoustic dynamics underlying expressive enactment. In this sense, the findings provide empirical support for the notion that musical emotion is embedded within coordinated action-sound patterns, measurable through large-scale oscillatory activity at the scalp level.

### Study limitations

The primary limitation of this work lies in its single-subject design, which constrains the generalizability of the findings. Single-participant EEG studies represent an established methodology in cognitive neuroscience when the goal is to characterize within-individual neural dynamics under carefully controlled conditions [36], and the present design was motivated by the need to preserve ecological validity during naturalistic, uninterrupted musical performance—a setting that would be difficult to standardize across multiple performers without compromising interpretative authenticity. Nevertheless, the idiosyncratic neural profile of a single individual, however expertly chosen, may not generalize to the broader population of concert pianists or trained musicians. Individual differences in cortical anatomy, motor strategy, and expressive style are known to produce substantial variability in EEG oscillatory patterns [19,20], and classification models trained on a single performer cannot be assumed to transfer to other individuals without retraining. The moderate dataset size (180 segments) represents a further constraint: while sufficient for exploratory analysis using classical machine learning under the present feature dimensionality, it limits statistical power and precludes robust evaluation of more complex architectures. The use of a relatively small held-out test set (36 segments across six classes) means that individual classification errors can substantially affect reported metrics, and results should be interpreted as indicative rather than definitive. Future work should prioritize multi-participant designs to disentangle performer-specific from general oscillatory patterns associated with expressive emotional states. Extension to additional performers would allow systematic examination of variability across expressive styles, motor strategies, and interpretative traditions, and would enable cross-subject generalization analyses that are currently not feasible. Larger datasets would also support the adoption of permutation-based significance testing and leave-one-out cross-validation schemes, providing more robust estimates of classification performance.

Future research may beneficially expand the feature space to include dynamic time-frequency representations, phase synchrony measures, and functional connectivity indices, which could capture additional aspects of neural coordination beyond static bandpower. In parallel, comparative evaluation of more advanced machine learning approaches, including deep learning architectures applied to larger datasets, may provide insight into hierarchical and nonlinear representations of affective-motor dynamics.

Finally, the present framework offers a promising foundation for future investigations into real-time decoding of expressive states during live performance. Such extensions could enable the development of adaptive musical interfaces, performer feedback systems, and interactive artistic technologies, bridging computational neuroscience and creative practice.

## Author Contributions

Conceptualization & data collection, A.M.P.; methodology, M.M; formal analysis, M.M.; investigation, A.M.P.; resources, A.M.P.; data curation, M.M. and A.M.P.; writing—original draft preparation, M.M & A.M.P. All authors have read and agreed to the published version of the manuscript.”

## Funding

This work was supported by Grant No. 2025-ATE-0077; project code 66232).

## Institutional Review Board Statement

The study was conducted in accordance with the Declara-tion of Helsinki and approved by the Local University Ethics Committee (CRIP, protocol number RM-2023-605) on 01/10/2023.

## Informed Consent Statement

Informed written consent was obtained from all subjects involved in the study.

## Data Availability Statement

All data supporting the findings of this study are included within the article. Additional information or data sharing are available from the author upon reasonable re-quest, particularly to support scientific collaboration.

## Acknowledgments

We are very grateful to Chang Qin for her support in stimulus validation.

## Conflicts of Interest

The authors declare no conflicts of interest.

## Disclaimer/Publisher’s Note

The statements, opinions and data contained in all publications are solely those of the individual author(s) and contributor(s) and not of MDPI and/or the editor(s). MDPI and/or the editor(s) disclaim responsibility for any injury to people or property resulting from any ideas, methods, instructions or products referred to in the content.

## Notes

### Competing Interest Statement

The authors have declared no competing interest.

